# Simultaneous multi-area recordings suggest that attention improves performance by reshaping stimulus representations

**DOI:** 10.1101/372888

**Authors:** Douglas A. Ruff, Marlene R. Cohen

**Affiliations:** Department of Neuroscience and Center for the Neural Basis of Cognition, University of Pittsburgh, Pittsburgh, Pennsylvania 15260

**Keywords:** *attention*, decoding, population analyses

## Abstract

Visual attention dramatically improves subjects’ ability to see and also modulates the responses of neurons in every known visual and oculomotor area, but whether those modulations can account for perceptual improvements remains unclear. We measured the relationship between populations of visual neurons, oculomotor neurons, and behavior during detection and discrimination tasks. We found that neither of the two prominent hypothesized neuronal mechanisms underlying attention (which concern changes in information coding and the way sensory information is read out) provide a satisfying account of the observed behavioral improvements. Instead, our results are more consistent with the novel hypothesis that attention reshapes the representation of attended stimuli to more effectively influence behavior. Our results suggest a path toward understanding the neural underpinnings of perception and cognition in health and disease by analyzing neuronal responses in ways that are constrained by behavior and interactions between brain areas.

## Introduction

Each of the huge number of psychophysical and physiological studies of visual attention show that attention profoundly affects subjects’ perceptual abilities and also modulates the responses of populations of neurons at every stage of visual and oculomotor processing^1–4^, Despite these oft replicated observations, whether any of the observed neuronal modulations can account for the improvements in psychophysical performance remains unknown. Two, non-mutually exclusive, hypotheses have dominated the literature (Figure 1A): that attention 1) improves visual information coding^5–7^, or 2) improves the efficiency with which visual information is read out by the premotor neurons involved in decision-making ^8–11^. The studies used to support these hypotheses were limited by available data and analysis methods, which primarily involved the responses of single neurons or pairs of simultaneously recorded neurons in the same brain area. We evaluated these hypotheses using the responses of groups of simultaneously recorded neurons in multiple stages of visuomotor processing, psychophysics, and data analysis methods that leverage that unique combination. We recorded simultaneously from groups of neurons in area MT, which encodes motion information ^12,13^ and the superior colliculus (SC), where neuronal responses are either visual, oculomotor, or intermediate, contribute to gaze control ^14–16^ and are involved in computing perceptual decisions ^17–19^. When we analyzed the responses of single neurons or pairs of neurons, we replicated previous observations, including the results from two of our previous studies, which focus on visual area V4 in two different tasks with spatial attention components: an orientation change detection task ^5^ and a contrast discrimination task ^6^. However, constraining our analyses of our MT data set or of both V4 data sets by the animals’ behavior and the simultaneous recordings from both areas made it clear that neither prior hypothesis constitutes a satisfying account of the observed attention-related improvements in performance.

**Figure 1.**
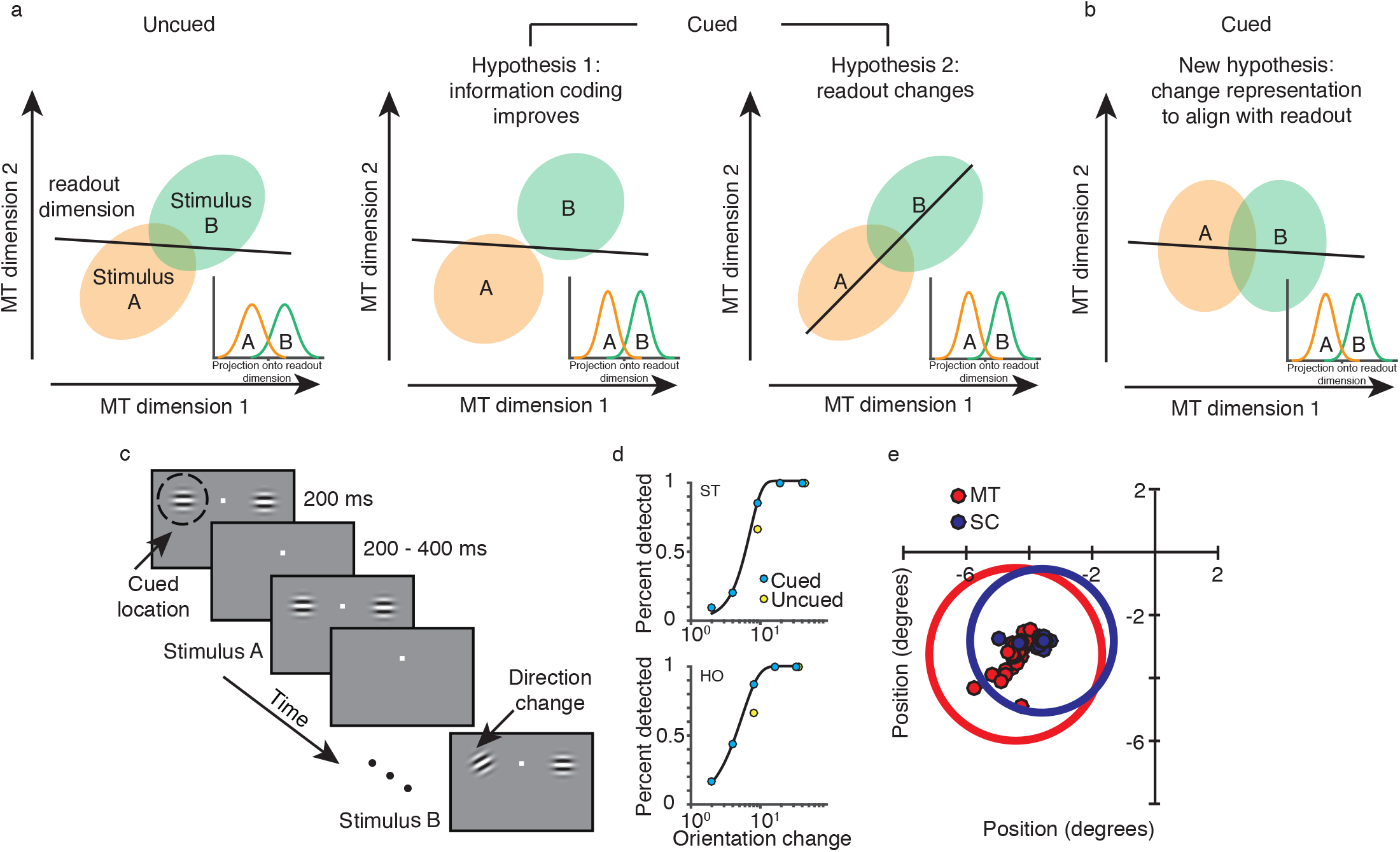
Hypotheses and methods. (A) Schematics describing predominant hypotheses about links between attention, visual cortical activity, and behavior. The left plot depicts MT population responses to two visual stimuli plotted along two dimensions in population response space (e.g. the first two principal components; see Methods) and a readout dimension which represents the visual information that is communicated to neuronal populations involved in planning behavior during the uncued condition. The insets depict projections of the population responses onto the readout dimension. Hypothesis 1 is that the MT representations of the two stimuli become more easily distinguishable (e.g. by separating the distributions of responses to the two stimuli). In this scenario, the distributions of projections along even a suboptimal readout axis may also be more separable. Hypothesis 2 suggests that attention changes the way visual information is read out from MT such that projections of MT population responses to the two stimuli onto the readout dimension are more separable. (B) Our new hypothesis: attention reshapes population responses so they are better aligned with relatively static readout dimensions. This alignment could be a direct result of widely observed attention-related changes in firing rates and response variability. (C) Direction change-detection task with cued attention. The drifting Gabor stimuli before the change were identical on every trial within an experimental session and can be thought of as stimulus A, while the changed stimulus can be thought of as stimulus B in the schematics in (A). (D) Psychometric curves from two example sessions (monkey ST, top, monkey HO, bottom) with best-fitting Weibull functions. Attention improved detection of median difficulty trials by 25% on average across all experiments (cued 76.5% detected across sessions, uncued 51.8% detected; N=15 sessions, two-tailed Wilcoxon signed-rank test, p=1.8×10^−4^). (E) Receptive field (RF) centers of recorded units from the same example session as in the top plot in (D). Dots represent the RF center (red, MT; blue, SC). The circle represents the size and location of the median RF from each area.

Our results suggest that on the timescale of perceptual decisions, across two visual areas and during both detection and discrimination tasks, spatial attention does not act primarily by improving information coding or by changing the way visual information is read out. Instead, the long-observed attention-related changes in the responses of visual cortical neurons account for perceptual improvements, but they do so by reshaping the representation of attended stimuli such that they more effectively drive downstream neurons and guide behavior (Figure 1B). Our study provides a framework for leveraging multi-neuron, multi-area recordings and controlled psychophysics to study how neuronal networks mediate flexible behavior in many systems, timescales, and tasks.

## Results

We compared evidence for and against two hypothesized attention mechanisms using neuronal responses collected while two rhesus monkeys performed the widely studied motion direction change-detection task in Figure 1C^5,9,20-22^, and then compared the results to recordings while monkeys performed a similar orientation change detection task ^5^ and a contrast discrimination task ^6^. As in the two previously published data sets, the animals’ performance in our new experiment was greatly affected (Figure 1D) by a cue instructing them to shift spatial attention between a stimulus within the same or opposite hemifield as the joint receptive fields of several dozen neurons that were recorded on multielectrode probes in MT (Figure 1E, red points) and the SC (blue points). MT and the SC represent different stages of perceptual decision-making and therefore provide the opportunity to evaluate each hypothesized attention mechanism. MT contributes to motion perception^12,13^. The SC is thought to play many roles in visually guided tasks including gaze control ^14–16^, decision-making ^17–19^ and attention ^4^.

### Population recordings replicate previously observed effects of attention

The two predominant attention hypotheses make different predictions about how attention should affect MT and the SC in our task. The first (information coding) hypothesis predicts that attention improves the motion direction information encoded in MT. The second (readout) hypothesis posits that attention changes the way that stimulus information is read out of MT to influence downstream responses and ultimately behavior. Our strategy was to show that our data are consistent with those in past studies by replicating the results that have been used as evidence to support each hypothesis and then to evaluate each hypothesis using analyses that leverage our simultaneous measurements from the subjects’ behavior and multi-neuron, multi-area recordings.

Past studies have evaluated these hypotheses by analyzing the responses of individual neurons or pairs of neurons, which typically lack the statistical power to reveal a strong link to behavior. Using our data set, we replicated the observations that have been used as evidence in favor of each hypothesis. Consistent with previous studies evaluating the information coding hypothesis ^2,3,23^, we found that attention increased the trial-averaged responses of neurons in both MT and the SC (Supplementary Figure 1A and B) and that attention decreased the extent to which the trial to trial fluctuations in neuronal responses to repeated presentations of the same stimulus are shared between pairs of MT neurons^5,7,21^ (quantified as the average spike count or noise correlation, or r_SC_^24^; Supplementary Figure 1C). Consistent with studies evaluating the readout hypothesis, attention increases correlated variability between the two areas^9,10,25^ (Supplementary Figure 1C). This attention-related increase was weakly dependent on the visual responsivity of SC neurons (Supplementary Figure 2).

The observed increase in correlations between areas suggests that attention-related effects are not simply due to global reductions in slow fluctuations, which has recently been hypothesized to explain attention-related correlation decreases within a single brain area^26,27^ (Supplementary Figure 3). On its face, this hypothesis seems unlikely to account for the spatially-specific effects of spatial attention (e.g. correlated variability increases in one hemisphere while decreasing in the other, even when neurons in the two hemispheres are simultaneously recorded^5^), meaning that reductions in the variability of global cognitive processes like arousal and motivation are unlikely to account for the attention-related changes in visual cortex. In addition to the observation that attention has opposite effects on noise correlations between pairs of neurons in the same than opposite areas, we found that attention has opposite effects on the local dynamics of the population responses within MT or the SC as it does on interactions between the two areas (Supplementary Figure 3C and 3D and Supplementary Figure 1C). These results are in conflict with the idea that the attention-related decrease in covariability within each area is a byproduct of a decrease in uncontrolled fluctuations in internal states, because such a decrease should, presumably, be brain-wide.

### Neuronal population decoding methods provide incomplete support the information coding or readout hypotheses

We reasoned that analyzing the relationship between populations of simultaneously recorded neurons in multiple brain areas with the animals’ behavior would provide insight into the relative importance of each hypothesized mechanism. To this end, we determined whether attention affects the amount of stimulus information that can be decoded from the population of MT neurons using cross-validated linear decoders that are optimized to a) dissociate between the original and changed stimuli (Stimulus decoder in Figure 2), b) predict the animals’ choices (whether or not they made an eye movement in response to change stimuli; Choice decoder), or c) predict the activity of the population of SC neurons we recorded (using responses to the original stimulus; SC decoder). These decoders were always constructed using data from trials with the intermediate change amount (see Figure 1D).

**Figure 2.**
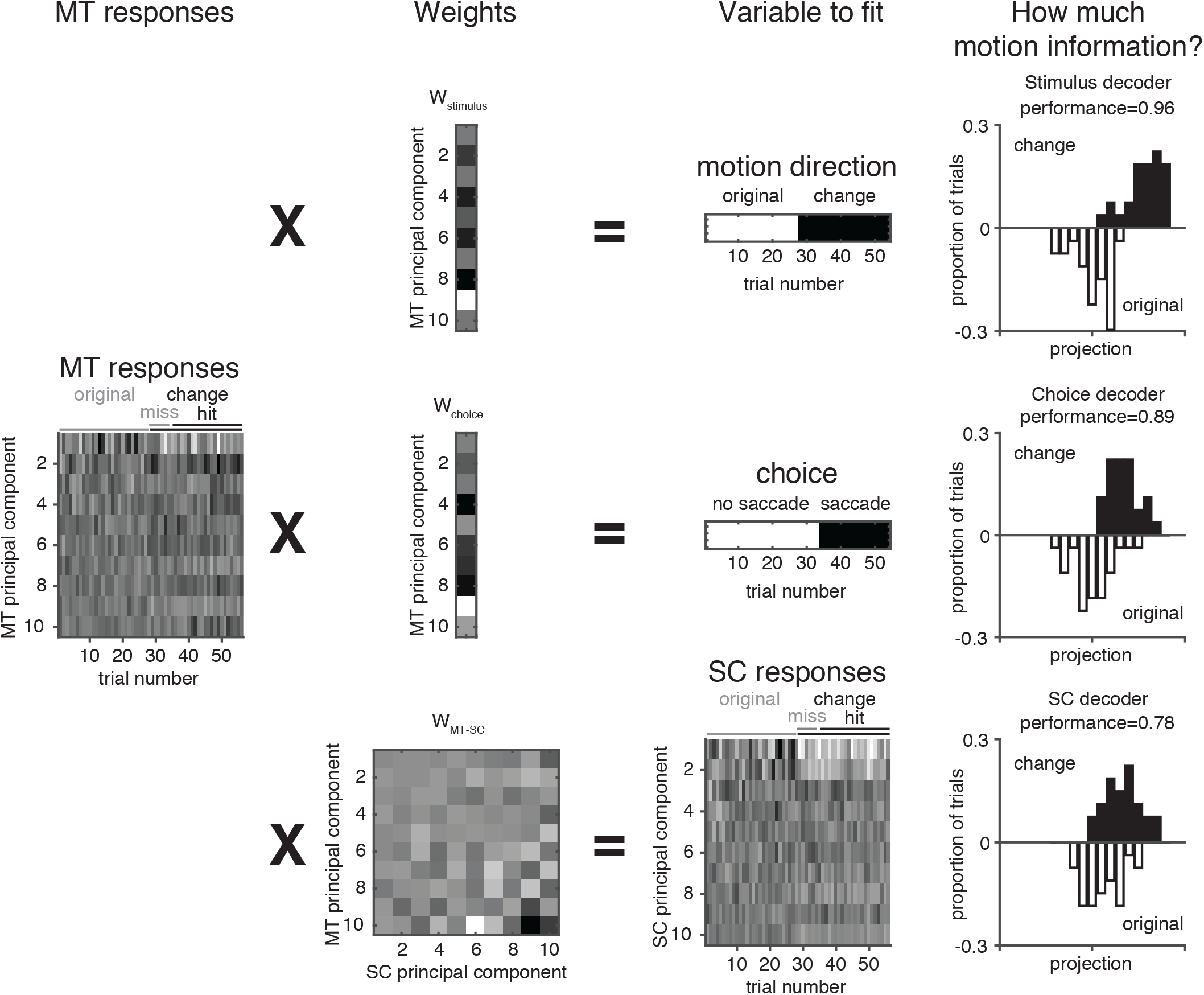
Schematic of our decoding procedure. We used linear regression to find the weights (second column) that best relate the first ten principal components of the MT population’s response (left) to both the original and change stimuli (Stimulus decoder; top row), the animal’s choice in response to change stimuli (Choice decoder; middle row), or the projections of the responses to the original stimulus of the population of simultaneously recorded SC neurons (SC decoder; bottom row). We assessed the performance of each decoder by decoding stimulus information from MT responses on a separate set of trials using each set of weights (right column) and responses to both the original and change stimuli. See methods for detailed decoding and cross validation procedures.

The information coding hypothesis posits that attention improves the stimulus information that could be gleaned by an optimal Stimulus decoder, but our data provided only weak support for this idea. Attention did not significantly affect the performance of an optimal decoder in our data set, even when we used a decoder optimized separately for each attention condition (Figure 3A, left bars). Recent theoretical work has demonstrated that high-dimensional decoders can ignore pairwise correlations that are orthogonal to the decoding axis and that correlations are increasingly likely to be orthogonal to this axis in larger populations ^28–30^. This suggests that the effects of attention on the stimulus information that can be decoded from small neuronal populations like the ones we recorded are likely to be even more minimal for larger populations, making it seem unlikely that attention-related improvements in information coding account for the robust improvements in behavioral performance that we observed.

**Figure 3.**
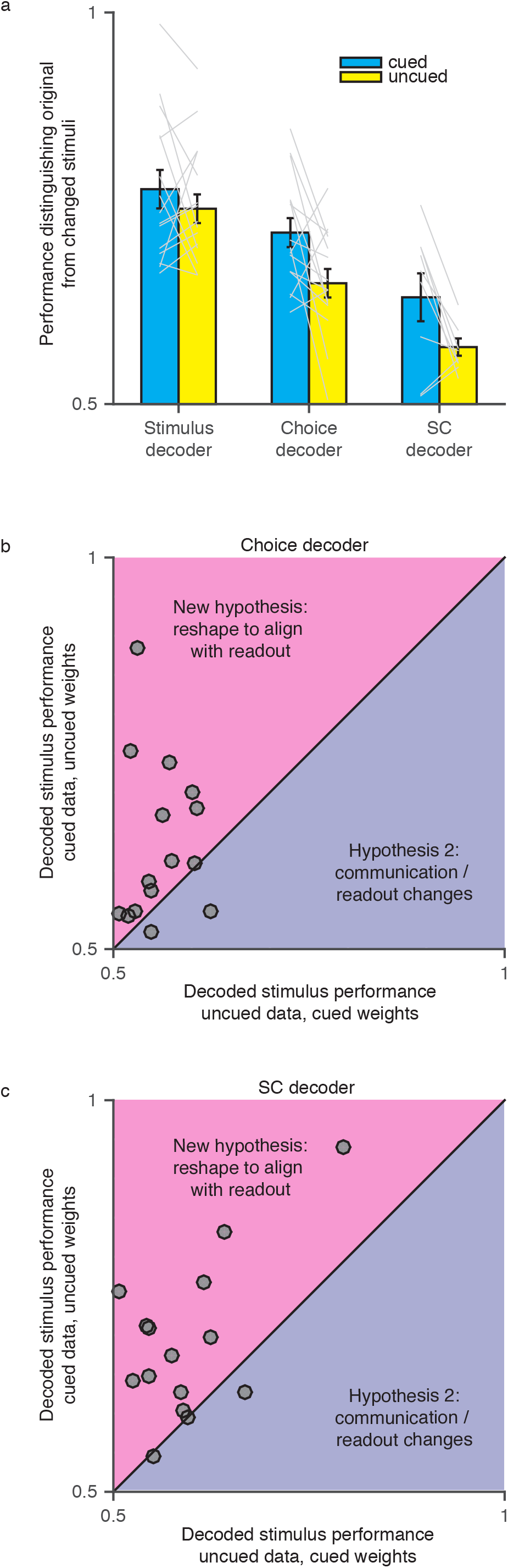
Effects of attention on the stimulus information that can be decoded from small populations of MT neurons. (A) Ability of a cross-validated linear decoder to distinguish the original from changed stimuli (intermediate change amount) for each decoder. Error bars represent SEM, gray lines are individual sessions. The effect of attention was significant for the Choice and SC decoders (N= 15 sessions, two-tailed paired t-tests, p=0.019 and p=0.048, respectively) but not for the Stimulus decoder (N= 15 sessions, two-tailed paired t-test, p=0.42). The effects of attention on the Choice and SC decoders were greater than for the Stimulus decoder (N= 15 sessions, two-tailed paired t-tests, p=0.023 and p=0.030, respectively), but not significantly different from each other (N= 15 sessions, two-tailed paired t-test, p=0.21). (B) Weight swapping analysis demonstrates that decoding performance was typically better using the MT responses from the cued condition and the Choice decoder weights from the uncued condition (y-axis) than using the MT responses from the uncued condition and the Choice decoder weights from the cued condition (x-axis; N= 15 sessions, two-tailed paired t-test, p=0.005). (C) Same, using the weights from the SC decoder (N= 15 sessions, two-tailed paired t-test, p=0.012).

The readout hypothesis posits that attention changes the importance of the attended stimulus in guiding behavior by changing the way its representation is read out by the neurons involved in computing decisions. Therefore, this hypothesis posits that attention should change the weights relating MT responses to either behavior or SC responses. We found that attention had larger effects on the stimulus information that is related to the animals’ choices on individual trials (Figure 3A, middle bars) or that is shared with the SC (Figure 3A, right bars) than it did on the Stimulus decoder. However, this difference could arise from either a weight change (Figure 1A) or a change within MT that results in more stimulus-related visual information being projected onto a static readout dimension (Figure 1B).

### A new hypothesis: attention reshapes sensory activity so that it more effectively guides decisions

Our data do not support the hypothesis that attention changes weights relating MT responses to SC responses or behavior. Because the responses of MT neurons are correlated and because the behavioral readout is binary, the weights obtained by each decoder are non-unique, making it impossible to identify weight changes by analyzing the weights themselves ^23,31^. However, we can infer their stability by measuring the stimulus information gleaned by each decoder using weights from the opposite attention condition from which they were calculated (see Methods). Both the Choice and SC decoders gleaned more stimulus information from MT responses in the attended than unattended condition when we used the weights computed in the opposite attention condition (Figures 3B and 3C). Together, these neuronal population analyses that use the animals’ behavior and the activity of downstream neurons to assess the hypothesized attention mechanisms reveal that neither the information coding nor readout hypothesis provide a satisfactory account of the large observed attention-related behavioral improvement.

Our observations suggest that in MT neurons recorded while monkeys performing a change detection task, attention acts primarily by changing the visual information that is used to guide behavior using relatively fixed readout weights. To investigate the generality of these observations to different visual areas and different tasks, we tested these hypotheses using two additional datasets. In the first dataset, monkeys performed the same direction change detection described here while we recorded from populations of V4 neurons^5^. Similar to our results in MT, we found that attention had larger effects on the stimulus information that is related to the animals’ choices (Choice decoder; Figure 4A) than it did on the stimulus information that could be gleaned using an optimal (Stimulus) decoder (Figure 4B). As in our MT data set (Figure 3B), the results from this data set suggest that attention typically reshapes V4 responses to align with relatively fixed readout mechanisms: decoding performance was typically better using the V4 responses from the cued condition and the Choice decoder weights from the uncued condition (y-axis) than using the V4 responses from the uncued condition and the Choice decoder weights from the cued condition.

**Figure 4.**
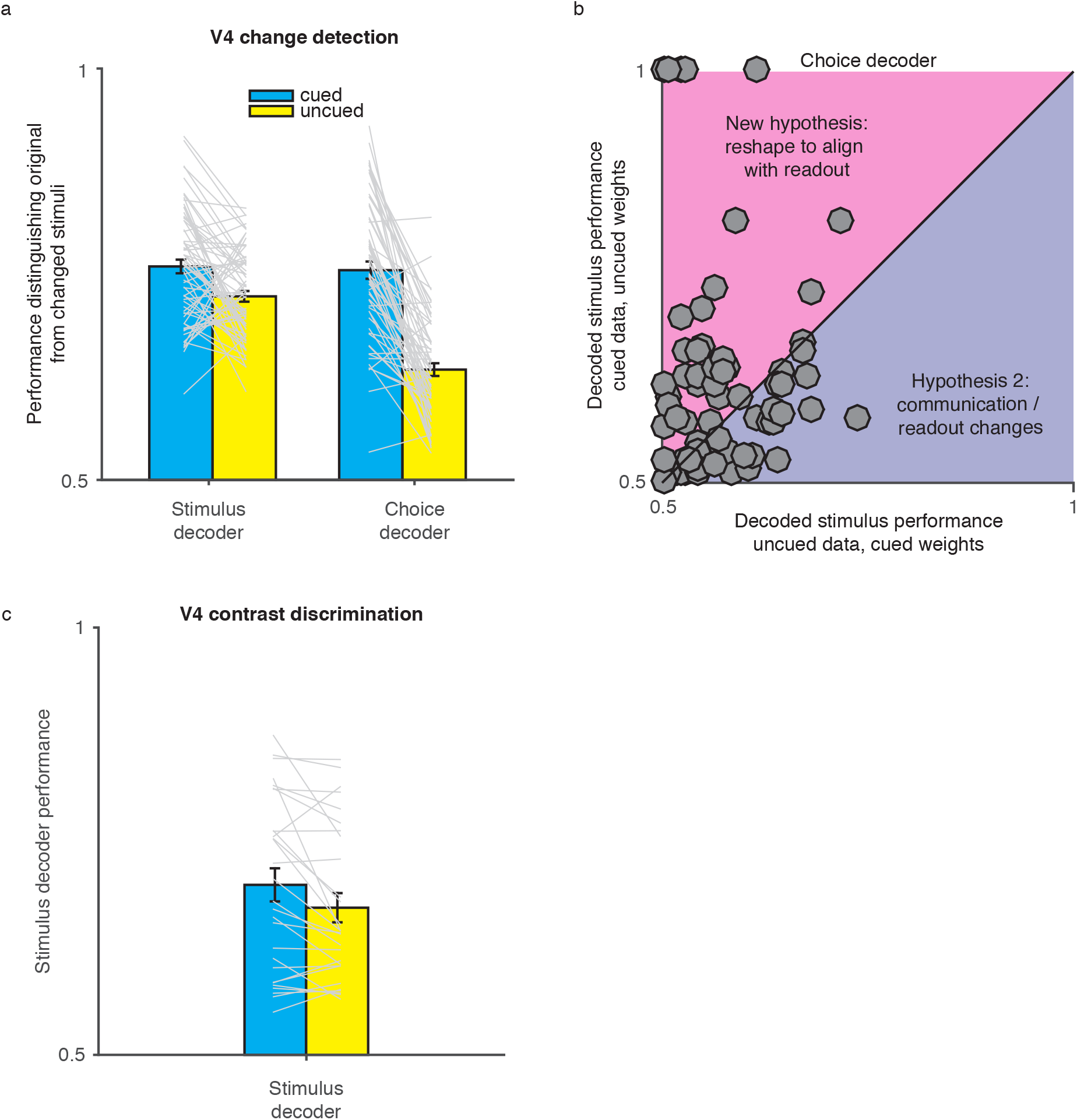
Similar attention-related effects on neuronal populations in two brain areas and two tasks. (A) In a change detection task, the effects of attention on the stimulus information that can be decoded from small populations of V4 neurons is similar to MT. The plot shows the ability of a cross-validated linear decoder to distinguish the original from changed stimuli (intermediate change amount) for both the Stimulus and Choice decoders (no SC data was available). Error bars represent SEM, gray lines are individual hemisphere-sessions (see methods). Attention significantly affected the performance of both the Stimulus and Choice decoders (N= 98 sessions, two-tailed paired t-tests, p=7.4×10^−4^ and p=1.1×10^−15^, respectively), but the attention-related improvement in the Choice decoder was greater than in the Stimulus decoder (N= 98 sessions, two-tailed paired paired t-test, p=2.1×10^−8^). (B) Decoding performance was typically better using the V4 responses from the cued condition and the Choice decoder weights from the uncued condition (y-axis) than using the V4 responses from the uncued condition and the Choice decoder weights from the cued condition (x-axis; N= 98 sessions, two-tailed paired t-test, p=0.0029; compare to Figure 3B). (C) The ability of a cross-validated linear decoder using V4 population responses to distinguish between stimulus configurations during a contrast discrimination task^6^ reveals no significant effect of attention (N= 17 sessions, two-tailed paired t-test, p=0.31). Plotting conventions as in A. Because of the details of the discrimination task (which did not include choices related to uncued stimuli), it was impossible to calculate a choice decoder using these data.

In the second new data set, we searched for attention-related changes in information coding in V4 neurons while monkeys performed a discrimination task^6^. These data provide a particularly important test of the information coding hypothesis because unlike in the change detection task in which attention has fairly uniform effects on V4 and MT neurons (increasing rates and decreasing noise correlations), we showed that in our discrimination task, attention can flexibly increase or decrease noise correlations in a way that is broadly consistent with improving information coding. Despite these findings, the results of our decoding analyses were similar for the detection and discrimination tasks, meaning that we did not find strong evidence that attention improves the amount of stimulus information that can be optimally extracted from a population of visual neurons in either task (Figure 4C). Together, these results provide evidence that in multiple visual areas and visually-guided tasks, attention acts primarily to reshape population activity so that more stimulus information is used to guide behavior using relatively fixed decision mechanisms.

Our data support the hypothesis that attention reshapes the representation of attended stimuli to more effectively guide behavior (Figure 1B). In this scenario, the critical changes are in visual cortex. However, this reshaping does not result in a large improvement in the stimulus information that can be gleaned by an optimal Stimulus decoder. Instead, the modulated neuronal activity in MT better aligns with the readout dimensions using relatively static weights.

How could a reshaping of the representation of an attended stimulus be implemented? The simplest mechanism would make use of the oft observed signatures of attention such as changes in firing rate gain^2,3,23^ or pairwise noise correlations^5-7,9,20-22,32-37^. We investigated the possibility that these simple response changes can account for the attention-related improvement in the stimulus information decoded using both the Choice and SC decoders in two stages. First, to verify the prediction of the weight-swapping analyses (Figures 3B and 3C), we constructed a single Choice decoder for both attention conditions (Figure 5A) and determined that it captured the attention-related improvement in decoded stimulus information (compare the blue and yellow bars in Figure 5B). Second, we used those same weights to decode stimulus information from population responses constructed using the mean rates from the uncued condition and the residuals from the cued condition (green bar). We found that simply using residuals (which incorporate both response variability that is private to each neuron and that which is shared between neurons) from the cued condition was enough to completely account for the attention-related improvement in decoded stimulus information in both the Choice (Figure 5B) and SC decoders (Figure 5C). These common decoders captured the attention-related improvement in decoded stimulus information and using residuals from the cued condition completely accounted for the attention-related improvement in decoded stimulus information.

**Figure 5.**
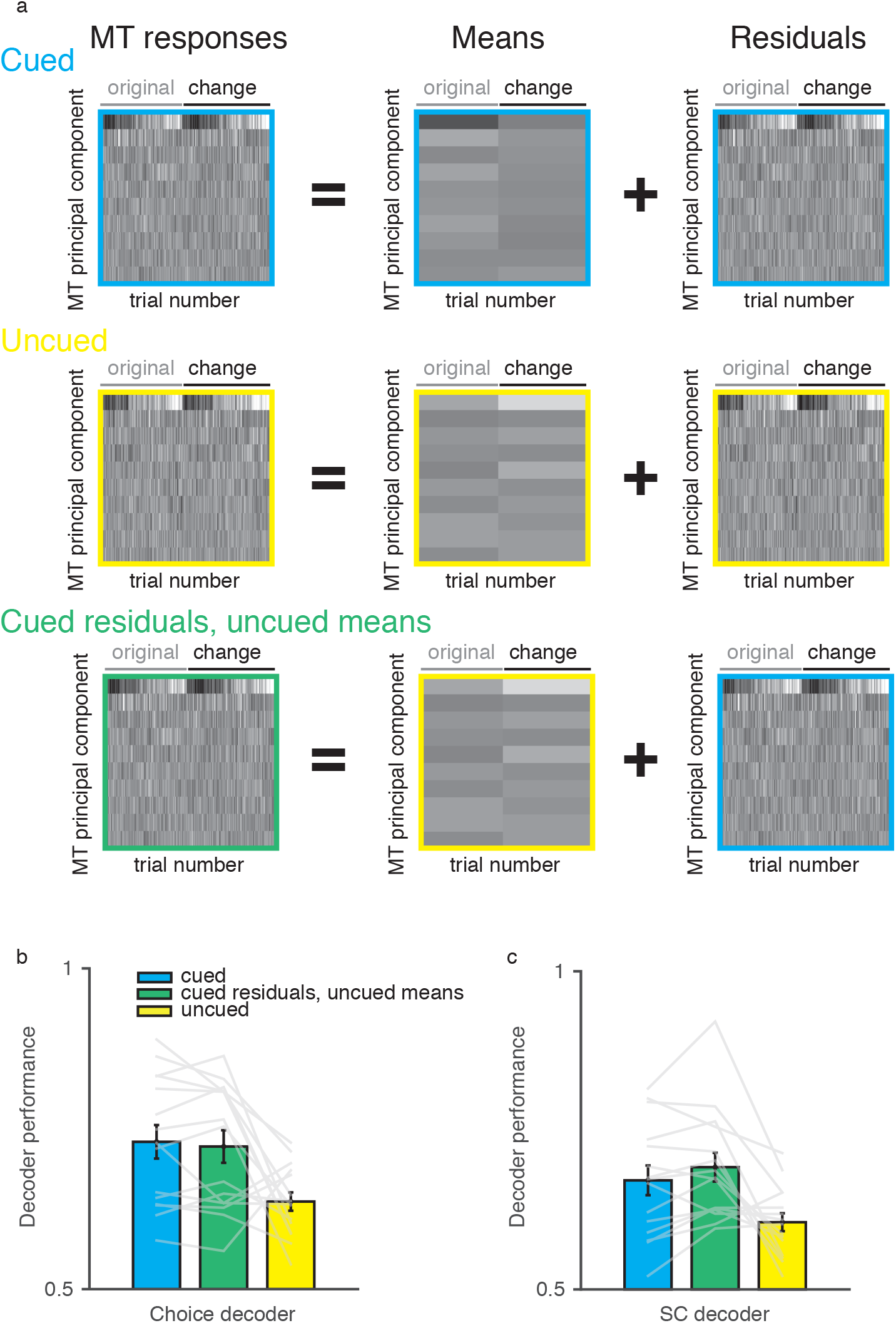
Effects of attention on the stimulus information that can be decoded from small populations of MT neurons is explained by changes in response variability. (A) Schematic of our procedure to understand which attention-related changes could account for the improvement in the amount of stimulus information that could be gleaned using the Choice decoder. We separated the first ten principal components of the MT population response (left) to the original and changed stimulus in both attention conditions into mean responses (scale adjusted to account for smaller value range) and residuals. We assessed the extent to which decoder performance was affected by attention-related changes in means and residuals by decoding stimulus information from MT responses on a separate set of trials in each attention condition and also using the residuals from the cued condition and the mean responses from the uncued condition (third row). See methods for detailed decoding and cross validation procedures. (B) Using the procedure described in (A), we found that the reshaping of the MT representation of the attended stimulus can be accomplished as a result of attention-related changes in response variability (e.g. noise correlations). The amount of stimulus information that can be decoded using a single Choice decoder whose weights are determined from data from both attention conditions is indistinguishable for the cued data and data constructed using the mean responses from the uncued condition and the residuals from the cued condition (N= 15 sessions, two-tailed paired t-test, p=0.84). Error bars represent SEM, gray lines are individual sessions. (C) Same as B, for the SC decoder. The amount of stimulus information that can be decoded using a single SC decoder whose weights are determined from data from both attention conditions is indistinguishable for the cued data and data constructed using the mean responses from the uncued condition and the residuals from the cued condition (N= 15 sessions, two-tailed paired t-test, p=0.48).

## Discussion

We used multi-neuron, multi-area recordings and psychophysics in detection and discrimination tasks to test two previous hypotheses and one novel hypothesis about the relationship between attention-related changes in perception and in neuronal responses on the timescale of perceptual decisions. In contrast with the hypotheses motivating most of the extensive literature concerning the neuronal basis of attention, our data are most consistent with the novel hypothesis that attention reshapes population activity so that information about the attended stimulus is read out to guide behavior. Our conclusions are based on comparing the visual information that can be gleaned from decoders optimized for the stimulus, the animals’ choices, and the activity of groups of visuomotor neurons. These results support the idea that behavioral flexibility is mediated by reshaping the representation of visual stimuli rather than improvements in information coding (which may be impossible given the immense amount of sensory information encoded in the brains of even anesthetized animals^30^ or in the responses of single neurons^13^) or by changing read out, which may be difficult to flexibly alter on the comparatively rapid timescale on which subjects can behaviorally shift attention^38^.

The idea of reshaping sensory information to better align with static read out mechanisms seems like it would require much more exotic mechanisms than the other hypothesized attentional mechanisms. However, we showed that commonly observed effects of attention on neuronal response variability were sufficient to reshape the representation of attended stimuli so that they more effectively influence the activity of downstream neurons and behavior (Figure 5B and 5C). Changing covariability may require a simpler mechanism than changing information coding or synaptic weights: we showed recently in a model that the covariability of a population of neurons can be readily changed by altering the balance of inhibition to excitation ^39,40^.

Although many studies are based on the implicit assumption that one or both of the information coding and communication hypotheses are true, there have been several recent studies that have failed to support the strongest versions of these hypotheses, and our reshaping hypothesis unifies these results. Mante and colleagues found the presence of both task-relevant and irrelevant information in prefrontal cortex, suggesting that the task, or attention-related gating of information does not occur in earlier stages of processing such as in visual cortex ^41^. This observation raises the question of why sensory responses are modulated if neurons near the end stages of processing in prefrontal cortex still encode task irrelevant information. To this point, Krauzlis and colleagues have suggested that attention-related changes in sensory cortex may arise as a byproduct of the process that interprets these signals ^42^. Further, a variety of experimental conditions that involve changing reward value ^43^ or saccade planning ^44^ result in changes in sensory responses that suggest a dependence on how the animal will use the sensory information. The reshaping hypothesis that we propose here is consistent with all of these findings, suggesting that sensory responses are modulated by the task such that the relevant information affects behavior and the irrelevant information is retained, perhaps for future actions or memory. Our findings suggest that this reshaping is achieved by changes to correlated variability early in visual processing, not by changing readout weights.

The idea that changing correlated variability better aligns sensory responses to a fixed readout is also consistent with our recent observation that in the change detection task, monkeys’ choices are well-aligned with the axis in population space that explains the most correlated noise ^21^. One exciting possibility is that the correlated variability axis represents the fixed readout dimension, perhaps because it is well-positioned to decode the motion direction of the broad set of stimuli that animals encounter outside the limited environment of most laboratory tasks ^23^. If so, reducing noise correlations and increasing firing rate gains would improve the stimulus information projected along that readout axis (following the intuitions in ^45^).

While our results were broadly consistent across two tasks and two visual cortical areas, it remains possible that attention uses different mechanisms in different tasks, brain areas, or sensory modalities. In particular, it is possible that the mechanisms underlying change detection, which is an important component of natural vision, are different than other tasks or that the mechanisms differ by brain areas. Therefore, the observation that attention also does not change the amount of stimulus information that can be decoded from visual cortex during a contrast discrimination task provides strong independent support for the generality of our findings.

However, even if we happened upon a special, albeit common, scenario using these two tasks, it is remarkable to observe a situation in which the large attention-related change in behavioral performance can be accomplished without changing information coding or weights between areas. In contrast, theoretical models and machine learning techniques often accomplish flexibility in computation almost solely by changing weights ^46–49^. Our results constitute an existence proof: an example of a situation in which flexibility can be mediated by simple changes within sensory cortex.

In the future, it will be interesting to use the same approach to determine whether similar mechanisms can account for behavioral changes associated with other cognitive processes (e.g. task switching) that might seem more likely to change the weights relating stimulus information to downstream neurons or behavior. Further, many neuropsychiatric disorders (including disorders of attention, Autism, and schizophrenia) are thought to involve changes in the same computations thought to underlie attention^50^. An exciting possibility is that these changes might be identified and potential therapies evaluated in animal models using the combination of behavioral evaluation and multi-neuron, multi-area recordings that we described here.

## Code availability

Custom Matlab code is available upon reasonable request to the authors.

## Data availability

The data that support the findings of this study are available from the corresponding author upon reasonable request.

### Acknowledgements

We are grateful to John Maunsell for granting permission for the use of the V4 data, to Karen McKracken for providing technical assistance, and to Faisal Baqai, Matt Getz, Jay Hennig, Chengcheng Huang, Amy Ni and John Maunsell for comments on an earlier version of this manuscript. Support to MRC from US NIH grants R00EY020844, R01EY022930, and Core Grant P30 EY008098s, the McKnight Foundation, the Whitehall Foundation, the Sloan Foundation, and the Simons Foundation.

## Author contributions

Both authors conceived and designed the experiments, analyzed the data and wrote the manuscript. D.A.R. collected the data.

## Online Methods

### Materials and Methods

The subjects of the simultaneously recorded MT and SC experiments were two adult male rhesus monkeys (*Macaca mulatta*, 8 and 9 kg and 8 and 6 years old, respectively). All animal procedures were approved by the Institutional Animal Care and Use Committees of the University of Pittsburgh and Carnegie Mellon University.

We presented visual stimuli using custom software (written in MATLAB using the Psychophysics Toolbox ^51,52^ on a CRT monitor (calibrated to linearize intensity; 1024×768 pixels; 120 Hz refresh rate) placed 54 cm from the animal. We monitored eye position using an infrared eye tracker (Eyelink 1000; SR Research) and recorded eye position and pupil diameter (1000 samples/s), neuronal responses (30,000 samples/s), and the signal from a photodiode to align neuronal responses to stimulus presentation times (30,000 samples/s) using hardware from Ripple.

### Behavioral Task

As previously described^5^, a trial began when the monkey fixated a small, central spot within a 1.25° per side square fixation window in the center of a video display while two peripheral full contrast, drifting Gabor stimuli (one overlapping the receptive fields of the recorded neurons, the other in the opposite visual hemifield) synchronously flashed on (for 200 ms) and off (for a randomized period between 200-400 ms) until, at a random, unsignaled time, the direction of one of the stimuli changed from that of the preceding stimuli (Figure 1C). The monkey received a liquid reward for making a saccade to the stimulus that changed within 450 ms of its onset. Attention was cued (using instruction trials prior to each block) in blocks of 50-100 trials, and randomly alternated between blocks where attention was cued to either the left or the right stimulus. In each block, the direction change occurred at the cued stimulus on 80% of trials, and at the uncued stimulus in 20% of trials (all uncued changes used either the middle or largest direction change, Figure 1D). In order to encourage fixation on longer trials, catch trials, in which no stimulus changed direction and monkeys were rewarded for maintaining fixation, were randomly intermixed throughout each block and made up approximately 12% of total trials. Psychometric data were fit with Weibull functions. Before recording commenced, the monkeys were extensively trained to have stable thresholds across a range of spatial locations (3-6 months). Because we recorded from several dozen neurons simultaneously, we could not optimize the stimuli for all neurons. We made sure to position one Gabor stimulus in the joint receptive field of the recorded neurons in both areas and we made an effort to set the properties of the size (approximately 3-6 degrees of visual angle), speed (approximately 3-12 degrees of visual angle per second) and direction of the stimuli so that they drove as many MT units as possible. The direction of all of the stimuli prior to the direction change (termed original stimulus) was constant throughout a recording session and this direction was typically either the median or mode of the distribution of MT preferred directions from that session. The range of direction changes differed from session to session, was selected based on the animals’ training history and depended on stimulus properties such as eccentricity and size. A typical range of change amounts for both animals was 1-35 degrees in log-spaced steps.

### Electrophysiological Recordings

Using linear 24 channel moveable probes (Plexon), we simultaneously recorded extracellular activity from direction-selective neurons in area MT and neurons in the superior colliculus that responded either visually, prior to a saccade, or both. Before beginning the experiment, we searched for neurons in both areas that had overlapping spatial receptive fields (Figure 1E) as determined by mapping with both drifting gratings and a delayed saccade task. The dataset consisted of a total of 306 responsive MT units and 345 responsive SC units total (36-58 units per session, mean 20 in MT, 24 in the SC for Monkey HO; 36-53 units per session, mean 21 in MT, 22 in SC for Monkey ST) in both MT and the SC in the right hemisphere using moveable, linear 24-channel V-probes (Plexon; inter-electrode spacing in MT = 50μm, SC = 100μm). We presented visual stimuli and tracked eye position as previously described^9^. The data presented are from 6 days of recording for Monkey HO and 9 days of recording for Monkey ST. Each day consisted of multiple blocks of the attention task (Figure 1C; mean 1015 of trials for Monkey HO, 745 for Monkey ST) preceded by receptive field mapping using a delayed saccade task and direction tuning during passive fixation.

### Data Analysis

All spike sorting was done offline manually using Offline Sorter (version 3.3.5; Plexon). We based our analyses on both single units and multiunit clusters and use the term “unit” to refer to either. Neuronal analyses in Supplemental Figure 1 and 2 used spike count responses between 50-250 ms after stimulus onset to account for visual latencies in the two areas. To remove response contamination from eye movements during change stimuli, data presented in the decoding analyses in Figure 3 and 4 used shorter response windows. Responses to both original and changed stimuli were measured from 50-185 ms after stimulus onset for monkey HO and 50-220 ms for monkey ST. These times were selected based on the distribution of each animal’s reaction times with the goal of maximizing the number of trials that could be included in the analyses. Trials with reaction times that began during those windows were excluded. Using these shorter response windows did not qualitatively affect the measures of attention described in Supplemental Figure 1. Attention still increased the firing rates of MT (mean attention index = 0.034, median attention index = 0.034; N = 306 units, two-tailed Wilcoxon signed rank test, p=1.2×10^−17^) and SC neurons (mean attention index = 0.071, median attention index = 0.05; N = 345 units, two-tailed Wilcoxon signed rank test, p=4.2×10^−44^) and decreased noise correlations within MT (N= 3285 pairs, two-tailed Wilcoxon signed rank test, p=7.7×10^−17^). To minimize the impact of adaptation on our results, we did not analyze the first stimulus presentation in each trial. We only analyzed a recorded MT unit if its stimulus-driven firing rate was 10% higher than its firing rate as measured in the 100 ms prior to the onset of the first stimulus. We only analyzed a recorded SC unit if its stimulus-driven firing rate was 10% higher than its firing rate as measured in the 100 ms prior to the onset of the first stimulus or if its response during a 100 ms epoch prior to a saccade on hit (correct) trials to the contralateral side was 10% larger than that same baseline. Stimulus presentations during which a microsaccade was detected were excluded from analyses^9,53^).

For firing rate analyses in Supplementary Figure 1A and B, attention indices were calculated using average spike counts on the (original) stimulus presentation prior to correct detections of the intermediate change amount depending on whether attention was directed into or out of the receptive fields of the recorded neurons using the formula (attend_in_ – attend_out_)/(attend_in_ + attend_out_). For illustrative purposes, significance of individual units was determined by a two-tailed paired t-test (p<0.05).

### Noise correlations

We defined the correlated variability of each pair of simultaneously recorded units (quantified as spike count correlation or r_SC_^24^) as the Pearson correlation coefficient between the responses of the two units to repeated presentations of the same stimulus. This measure of r_SC_ represents noise correlations rather than signal correlations because the responses used in this analysis were always to an identical visual stimulus. For Supplementary Figure 1C, we included responses from stimulus presentations 2 though 10 from trials that ended with either a hit, miss or correct catch trial and that were immediately followed by the maintenance of fixation and continuation of the trial (i.e., stimulus presentations where the behavioral response on the subsequent stimulus presentation was not a saccade). We z-scored responses as a function of the stimulus presentation number in each trial and then pooled data across stimulus presentations before calculating noise correlations. Results did not qualitatively change if we did not perform this z-score procedure. For Supplementary Figure 1D, we included data from all stimulus presentations prior to the change stimulus (except the first) and sorted them depending on what the behavioral outcome was on the subsequent stimulus presentation. Pairs of units that were recorded on the same electrode were not included in correlation analyses. The data presented in Supplementary Figures 1C consisted of 3,285 MT pairs, 3,948 SC pairs and 6,934 between area pairs.

### Decoding

We focused our decoding analyses (Figures 2, 3 and 5) on trials in which the third largest (middle) direction change occurred, because changes of that magnitude occurred in both attention conditions. This approach also serves to linearize the problem by attempting to classify between one of two directions of motion. Therefore, we have restricted our decoding approach to using linear methods. We performed the decoding analyses using responses from trials that were either hits (correct detection) or misses (maintained fixation after change stimulus). All of the data sets contained at least 10 trials in each attention condition and at least three hits and three misses in each condition. We did not include false alarms in the analyses because there were too few (and they were too inconsistent across recording sessions) to handle appropriately.

We used the decoding strategy schematized in Figure 2. We began by constructing a matrix of MT responses for each attention condition: ‘MT responses’ (a # MT neurons by 2*# trials matrix of MT responses to the stimuli before the direction change and the changed stimulus on the relevant trials). The stimulus decoder was performed using two matrices for each attention condition: all of ‘MT responses’ (a # MT neurons by 2*# trials matrix of MT responses to the stimuli before the direction change and the changed stimulus on the relevant trials) and ‘motion direction’ (a 1 by 2*# trials vector of zeros for the stimulus before the change, referred to as ‘original’, and ones for the changed stimulus, referred to as ‘change’). The Choice decoder was performed using two matrices for each attention condition: the responses during change stimulus presentations from ‘MT responses’ (a # MT neurons by 1*# trials matrix of MT responses to the change stimulus on the relevant trials) and ‘choice’ (a 1 by 1*# trials vector of zeros for change stimulus presentations on which the animal did not make an eye movement, referred to as ‘no saccade’, and ones when the animal made an eye movement, referred to as ‘saccade’). The SC decoder was performed using two matrices for each attention condition: the responses during original stimulus presentations ‘MT responses’ (a # MT neurons by 1*# trials matrix of MT responses to the original stimulus on the relevant trials) and ‘SC responses’ (a # SC neurons by 1*# trials matrix of SC responses to the original stimuli on the relevant trials). We refer to this final decoder as an ‘SC decoder’ but the weights are defined with no directionality: we have simply identified the weights that best relate the activity between the two areas. We used only responses to the original stimulus for the SC decoder because of the strong presaccadic responses present during changed stimuli.

We cross validated by holding out the two stimulus presentations from ‘MT responses’ (for the original and changed stimuli) from one trial at a time to perform the classification of motion direction. To reduce the number of weights we needed to fit and therefore improve our confidence in the weights we did fit, we performed PCA on the MT and SC responses to find the first 10 PCs in each area. The choice of number of vectors did not qualitatively affect the results in the range of 4-15 vectors. We then performed linear regression to find the weight vectors (for the Stimulus and Choice decoders) or weight matrices (for the SC decoder) that related projections along the first ten MT PCs plus a vector of ones to ‘motion direction’, ‘choice’, or projections along the first 10 SC PCs in each attention condition.

We assessed the stimulus information in each decoder (Figure 3) by multiplying projections of MT responses to the original and changed stimuli from the held-out trial by the fitted weights and either determining whether those weighted sums correctly classified the stimuli as original or changed (Stimulus and Choice decoders) or whether a linear classifier correctly classified those stimulus presentations on the basis of the predicted SC responses (SC decoder). The performance of the decoder is defined as the area under the receiver operating characteristic curve comparing the distributions of weighted average responses to each stimulus using the weights constructed for each decoder.

The critical aspect of the decoding analysis is that we ask how much *stimulus* information is contained in each different subset of MT activity. The Stimulus (or optimal) decoder will perform best, because it was designed specifically to ask this question. The Choice and SC decoders identify different subspaces of MT activity and then ask how much stimulus information is contained in those subspaces. These decoders, by definition, will perform worse than the Stimulus decoder, but they are asking the same question.

To assess the stability of the weights for each decoder in the two attention conditions, we assessed the stimulus information gleaned by each decoder using the sensory responses from one attention conditions and the weights calculated from the other (Figures 3 and 4). Because the responses of visual neurons are non-unique and because our behavioral response is binary, the weights found with our linear decoding methods are non-unique^23,31^. It is therefore not informative to make direct comparisons of the weights across conditions. Instead, we borrowed the spirit of the analyses in a recent study^31^ and compared the stimulus information that could be gleaned using each set of weights in each attention condition. In general, the choice and SC decoders performed better with weights computed from the same attention condition, even though we cross-validated these analyses (this effect could be attributed to non-stationarities in the recordings or the monkey’s behavior). The critical comparison is the performance of the decoders using sensory responses from one attention condition and weights from the other (Figures 3 and 4).

For the decoding analysis in Figure 5, we took a similar approach to the previously described Choice and SC decoders, except that we combined data from both the cued and uncued conditions to calculate decoding weights. We then decomposed the responses of the population responses to each stimulus in each attention conditions into mean responses and residuals (R=M+S, where R is the number of neurons by number of trials matrix of spike count responses to one stimulus in one attention condition, M is a matrix of mean responses for each neuron, and S is the matrix of residuals). We tested the hypothesis that attention-related changes in the residuals account for the improvement in stimulus information used to guide behavior by decoding stimulus information from responses created by using the mean responses from the uncued condition and residuals from the cued condition.

The analyses of the V4 data from the change detection task (Figure 4A and 4B) were identical manner to the MT data described above. This dataset consisted of multineuron recordings using Utah arrays placed in both hemispheres of V4 during 37 experimental sessions in two animals, the details of which are described in^5^. Data from each hemisphere was treated separately in the decoding analyses, so each session contributes two data points for each analysis (gray lines in Figure 4A). The details of the contrast discrimination task used in Figure 4C required a different form of the Stimulus decoder. This dataset consisted of multineuron recordings using Utah arrays placed in both hemispheres of V4 during 17 experimental sessions in two animals. The details of this experiment have been previously described^6^. Briefly, two monkeys judged which of two stimuli in a pair was higher contrast by making a saccade to a target representing its choice. Attention toward one pair of stimuli or the other was changed in blocks. The Stimulus decoder (Figure 4C) compares performance using V4 responses to distinguish between a given stimulus configuration and its opposite configuration in the attended and unattended conditions. As in the other V4 data set, data from each hemisphere was treated separately.

### Statistics

Paired tests, either two-tailed t-tests or non-parametric Wilcoxon tests, were employed for all statistical analyses. In cases where t-tests were used, the data distribution was assumed to be normal but this was not formally tested. No statistical methods were used to pre-determine sample sizes but our sample sizes are similar to those reported in previous publications ^6,9^. There was no way to perform data collection and analysis blind to the conditions of the experiments because our data were not grouped. Please see the Life Sciences Reporting Summary for additional information.

