## Supplemental Figures for "Simultaneous multi-area recordings suggest that attention improves performance by reshaping stimulus representations"

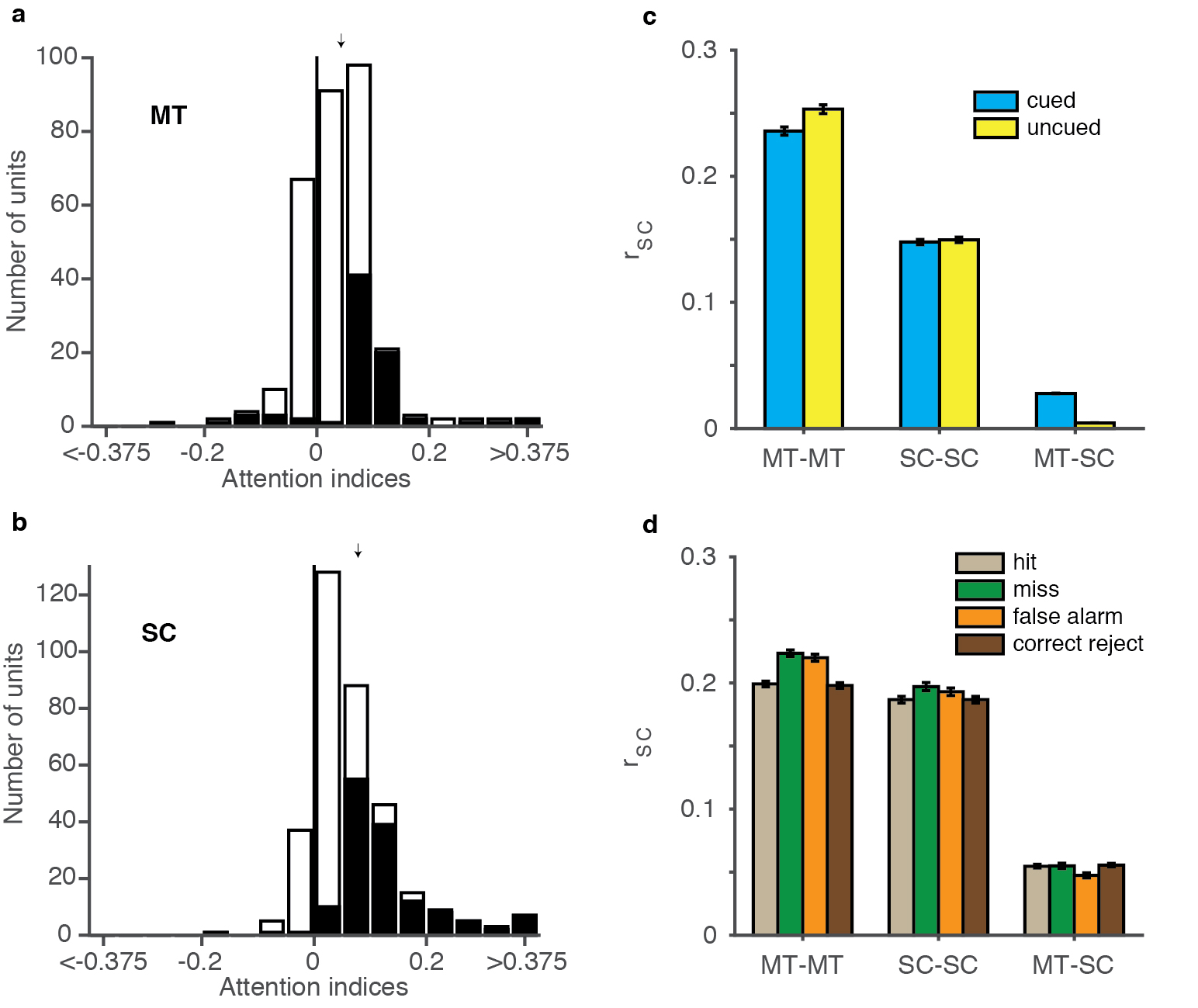

Supplementary Figure 1

Effects of attention on common analyses of individual units and pairs of units

(A) Attention increases firing rates in MT, quantified as the difference in firing rates in the different attention conditions divided by the sum. Units with significant differences in average responses for the two conditions are specified by black bars (two-sided t-test, p<.05). This distribution (mean = 0.04, median = 0.04) is significantly different from zero (N=306 units, two-sided Wilcoxon signed rank test, p=4.0X10^-22^). (B) Same as A, for SC data. This distribution (mean = 0.073, median = 0.05) is significantly different from zero (N=345 units, two-sided Wilcoxon signed rank test, p=4.8X10^-44^). (C) Within and between area noise correlations calculated from spike counts during stimulus presentations that preceded successful maintenance of fixation from trials that ended with either a hit or miss or were a successful catch trial. Attention decreases average correlations within MT (N=3,285 pairs, two-sided Wilcoxon signed rank test, p=6.6X10^-15^), not in the SC (N=3,948 pairs, two-sided Wilcoxon signed rank test, p=0.38) and increases them between the two areas (N=6,934 pairs, two-sided Wilcoxon signed rank test, p=5.7X10^-58^). Error bars are standard error of the mean. (D) Within and between area noise correlations calculated from spike counts that immediately preceded different behavioral outcomes during cued trials. Misses and false alarms are associated with higher correlations within MT (two-tailed t-test on hits and correct rejects versus misses and false alarms, p=0.0094) and SC (two-sided t-test on hits and correct rejects versus misses and false alarms, p=0.03) but not between the two areas (two-tailed t-test, p=0.23). Error bars are standard error of the mean.

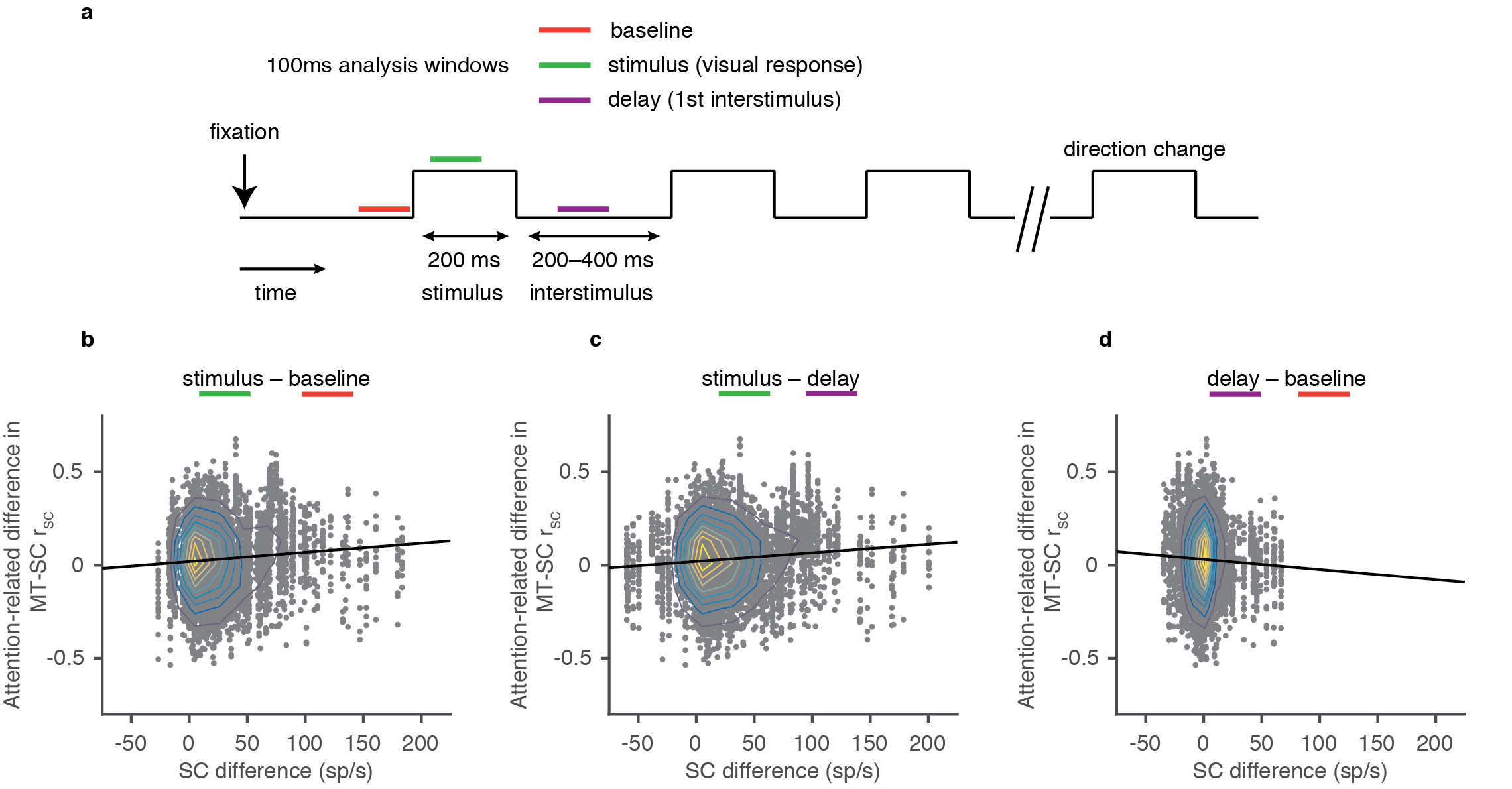

**Supplementary Figure 2**

Relationship between SC responses during different task epochs and attention-related correlation changes with MT.

(A) Schematic of task timing depicts the three 100ms epochs used to count spikes in SC units. The baseline period began 100ms before the first stimulus appeared, which is after stable fixation had been acquired. The stimulus period was shifted 30 ms after the appearance of the visual stimulus, to account for the earliest visual latencies observed in the SC. The delay period began 100ms after the first stimulus turned off and always ended prior to the onset of the second stimulus. (B) Attention-related changes in MT-SC r_SC_ plotted against the difference between each SC unit’s response during the stimulus and baseline periods. There are multiple MT-SC correlation differences measured for each SC unit. Correlations between MT and SC were calculated using the same data and methods as Supplementary Figure 1C (N=6,934 pairs, Pearson correlation, rho=0.088, two-sided t-test, p=2.4X10^-13^). Isolines depicting the decile boundaries are overlaid over the individual data points. (C) Similar to B, but data are now sorted by the difference between each SC unit’s response during the stimulus and delay periods (N=6,934 pairs, Pearson correlation, rho=0.091, two-sided t-test, p=2.8X10^-14^). (D) Similar to B, but data are now sorted by the difference between each SC unit’s response during the delay and baseline periods (N=6,934 pairs, Pearson correlation, rho=-0.037, two-sided t-test, p=0.0023).

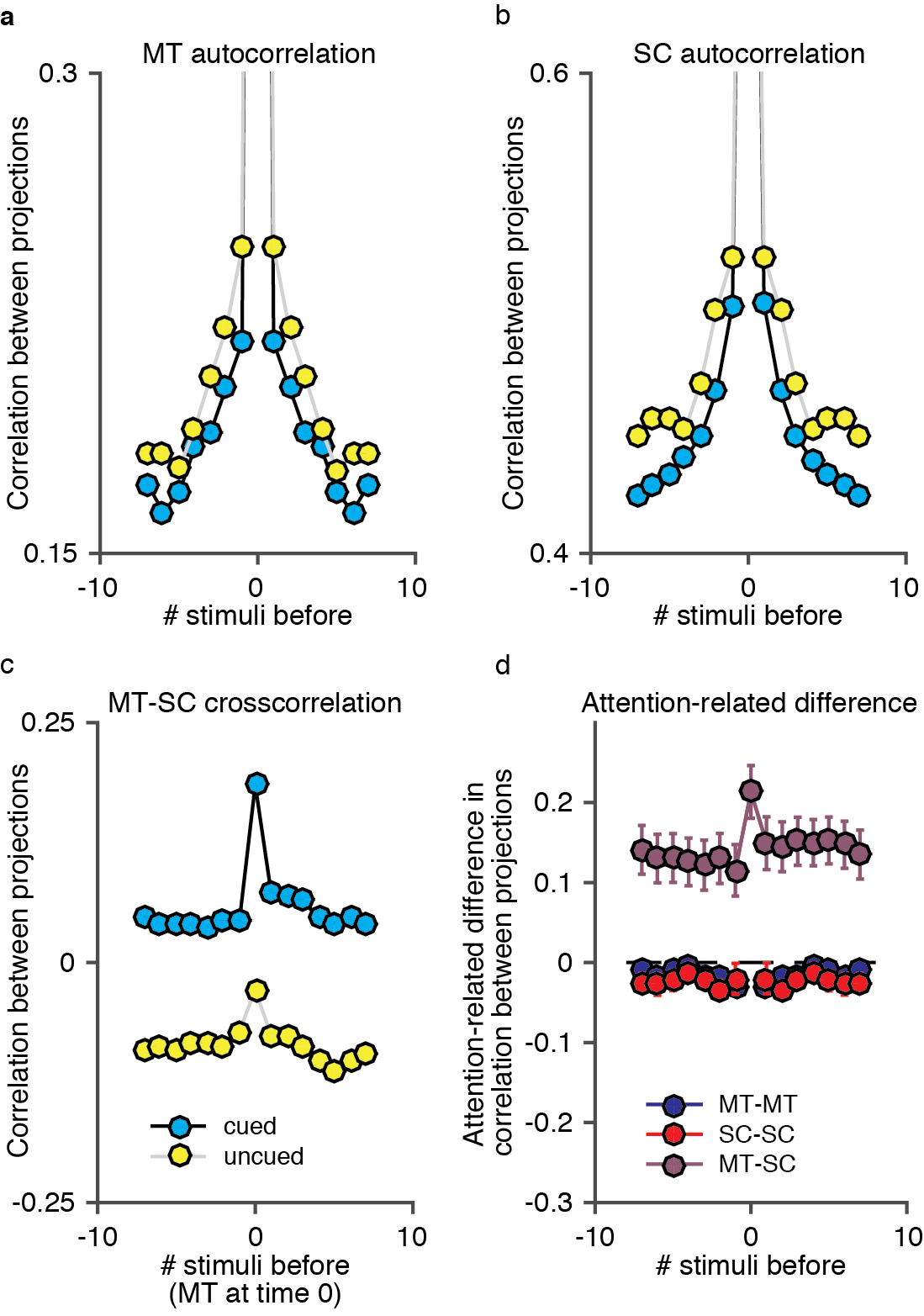

**Supplementary Figure 3**

Attention has opposite effects on slow fluctuations in neuronal population responses within and across areas.

(A, B) Autocorrelations between projections onto the first principal component of population responses to repeated presentations of the same visual stimulus in (A) MT, and (B) the SC. The x-axis plots time lag in units of stimulus presentations (400-600 ms). Because these were responses to identical stimuli, the first PC is the dimension in population space that accounts for the greatest shared variability across the population. (C) Cross correlation between projections onto the first PCs in MT and the SC (same data and plotting conventions as in A and B). (D) Attention-related difference in autocorrelation or cross correlations between the projections in the previous plots. Error bars represent standard error of the mean. Attention was associated with a statistically significant decrease in autocorrelation overall (t-tests, p=0.04 in MT and 0.02 in SC) in both areas and in 11/15 individual MT data sets and 9/15 SC data sets (bootstrap tests, p<0.05 with a Bonferroni correction) and a significant increase in cross correlation overall (t-test, p=0.0009) and in 11/15 individual data sets (bootstrap tests, p<0.05 with a Bonferroni correction).
